# Gastruloid development competence discriminates different states of pluripotency between naïve and primed

**DOI:** 10.1101/664920

**Authors:** Federica Cermola, Cristina D’Aniello, Rosarita Tatè, Dario De Cesare, Alfonso Martinez-Arias, Gabriella Minchiotti, Eduardo Jorge Patriarca

## Abstract

Floating spheroidal aggregates (aggregomes) of mouse embryonic stem cells (mESCs) can develop into polarized/elongated organoids, namely gastruloids. Here we report a high-performing assay to measure gastruloids formation efficiency (GFE), *i.e.* the fraction of gastruloid-developing aggregomes. By exploiting this procedure, we provide morphological and molecular evidence that gastruloid development relies on *Cripto*. We also demonstrate that GFE decreases as pluripotency progresses from naïve to primed state. Indeed, naïve ESC-derived aggregomes efficiently elongate (GFE≥95%), while primed EpiSCs fail to aggregate and consequently to generate gastruloids (GFE=0%). Conversely, while early-primed EpiLCs properly aggregate, EpiLC-derived aggregomes are mostly abortive (GFE=0%). Unlike EpiLCs, L-Proline-treated ESCs (PiCs) generate productive aggregomes (GFE≥50%), which however begin to elongate earlier and generate smaller gastruloids that appear more differentiated. Like EpiLCs, PiCs are competent to differentiate into primordial germ cell-like cells (PGCLCs), suggesting that PiCs capture an EpiLC-like state with unique competence for both gastruloid formation and differentiation into PGCLCs. Thus we propose GFE assay as a simple and robust *in vitro* method to discriminate different phenotypic/functional states of the pluripotency continuum.

## INTRODUCTION

Pluripotent stem cells (PSCs) forced to grow under non-adherent culture conditions are inclined to establish cell-cell adhesive interactions and, eventually, to generate floating three-dimensional cells aggregates (hereafter called ‘aggregomes’). Tightly packed and globular aggregomes of mouse embryonic stem cells (mESCs) can develop, under the pressure of specific cues, into organoids that break symmetry and grow in a polarized manner with regard to three orthogonal axes (Baillie-Johnson et al., 2015; Marikawa et al., 2009; Simunovic and Brivanlou, 2017; Turner et al., 2016; Turner et al., 2017; van den Brink et al., 2014; Vianello and Lutolf, 2019). These structures are called ‘gastruloids’ (van den Brink et al., 2014) and the *aggregome to gastruloid* transition has been shown to exhibit some features of mouse embryo gastrulation (Beccari et al., 2018; Turner et al., 2017). It has been found that gastruloid development depends on: i) the mESC line examined; ii) the diameter (150-200 μm) of the initial aggregomes, which is determined by the number of cells aggregated (200-400 cells); and iii) a transient (24 h) treatment of developing aggregomes with a WNT (<u>W</u>ingless/I<u>nt</u>egrated) signalling inducer (Baillie-Johnson et al., 2015; Turner et al., 2017). The process can be variable and give rise to atypical organoids including large spherical cell masses (proliferation without symmetry breaking), spherical structures with protrusions (multiple elongation attempts), and elongated gastruloid-like organoids but with one or few short ectopic protrusions (Baillie-Johnson et al., 2015; Turner et al., 2017). The reason for these failures remains largely unknown.

Gastruloid formation assays are performed using *in vitro* cultured PSCs, and it has been well established that PSC culture conditions can alter the grow behaviour of stem cells (Martello and Smith, 2014; Mulas et al., 2019; Smith, 2017). For instance, mESCs grown in 2i_LIF medium, *i.e.* a medium supplemented with a combination of MEK inhibitor PD0325901, GSK3 inhibitor CHIR99021 and Leukemia Inhibitory Factor (LIF), acquire a naïve/ground state of pluripotency (Ying et al., 2008). Under low-density 2D culture conditions, naïve cells generate tightly packed domed-shaped colonies. Conversely, when treated with a combination of Fibroblast Growth Factor and Activin A (F/A), following LIF withdrawal, mESCs acquire a primed state of pluripotency (Brons et al., 2007; Najm et al., 2011; Tesar et al., 2007). Certainly, after 2 days of treatment (2d_F/A), mESCs acquire features of germ-line competent pre-implantation epiblast-like state (early-primed epiblast-like cells, EpiLCs), whereas after 5 days of treatment (5d_F/A) the cells acquire features of the post-implantation epiblast (epiblast stem cells, EpiSCs). Notably, while the 2D colony phenotype of EpiLCs is unknown since they cannot be propagated, EpiSCs can proliferate indefinitely generating irregular flat-shaped colonies. The colony phenotypes suggest that while naïve cells are inclined to establish robust cell-cell adhesive interactions, EpiSCs are more prone to establish cell-substrate than cell-cell interactions. Besides the extreme naïve and primed states, intermediate states of pluripotency can be captured modulating the level of metabolites, such as L-Proline and Ascorbic acid (Vitamin C, VitC), supplied in the growth medium (Comes et al., 2013; D’Aniello et al., 2017a; D’Aniello et al., 2017b). Certainly, a high L-Proline regimen drives mESCs towards an early primed pluripotent state, which at a molecular level resemble that of EpiLCs. Notably, under 2D culture conditions, L-Proline-induced cells (PiCs) grow generating structurally complex colonies displaying a naïve-like domed-shaped centre surrounded by an irregular flat-shaped EpiSC-like monolayer of cells that progresses to a crown of free motile mesenchymal-like cells (Casalino et al., 2011; Comes et al., 2013; D’Aniello et al., 2019). In correlation, the cell-cell adhesion molecule E-cadherin is delocalised to the Golgi and focal adhesion-related genes are significantly deregulated in PiCs (Comes et al., 2013). These data support the idea that the naïve to primed transition is associated with a loss of the stem cell propensity to establish cell-cell interactions.

Based on these observations we reasoned that the pluripotency state could influence gastruloid formation efficiency (hereafter GFE); namely, the fraction of initial aggregomes that becomes a fully developed gastruloid. Our results lead us to propose GFE assay as a robust *in vitro* method to discriminate different states of pluripotency.

## RESULTS

### Improvement of Gastruloid Formation Efficiency (GFE) in mESCs

In order to optimise the efficiency of gastruloid formation, we have adapted the current protocol of Baillie-Johnson et al., 2015, using a TBV2 mESC line (Casalino et al., 2011) with a particular emphasis on increasing ESC counting accuracy and precision. We first verified the ability of TBV2 mESCs to generate gastruloids using the protocol described by Baillie-Johnson et al., 2015. To this end, FBS/LIF mESCs were seeded at the appropriate density (300 cells/40 μl) in ultra-low attachment plates to force aggregation. Forty-eight hours after aggregation (AA), we observed the formation of spherically shaped cell aggregomes with a diameter ranging from 125 to 195 μm (mean = 156 μm) (**Fig. S1A**). Three days later (120h AA), a major fraction (~75%) of the primary cell aggregomes had become elongated-shaped gastruloids (0,5 to 1 mm long), whereas a minor but significant fraction remained as unstructured globular cell masses (~10%) or developed into abnormal/aberrant organoids (~15%) displaying one or more ectopic elongation zones/protrusions (**Fig. S1B-C**). These results confirmed the occurrence of frequent developmental failures during gastruloid formation assay, as already observed with different mESC lines analysed (Baillie-Johnson et al., 2015; Turner et al., 2017).

In an attempt to improve the gastruloid formation efficiency (GFE), and specifically to minimize the fraction of abnormal/abortive events, we modified the culture conditions, as well as the dissociation and seeding procedures to induce cell aggregation (see **Fig. 1A**). First, since the morphology of the cell colonies reflect their pluripotency state (**Fig. S1D**), we used cells grown at low density rather than as confluent (>60%) monolayer. Thus, to perform GFE assay, confluent FBS/LIF TBV2 mESCs were first dissociated by trypsin digestion and then seeded at low density (250 cells/cm^2^) on gelatin-coated plates in N2B27 supplemented with LIF and the 2i inhibitors (PD0325901/CHIR99021) (naïve state-inducing medium). Under this culture conditions, mESCs gave rise to 90-95% of naïve/round-domed cell colonies (**Fig. S1D**). Second, given that cell dissociation with trypsin reduces cell-cell adhesion capability of early primed (*i.e.* L-Pro-induced; Comes et al., 2013) and primed (F/A treated) mouse pluripotent stem cells (Brons et al., 2007), which may affect the subsequent aggregation step and eventually reduce the GFE, cells were dissociated with a milder accutase mix treatment (**Fig. 1A**). Third, since proper gastruloid development relies on the number of aggregated cells (Baillie-Johnson et al., 2015), a precise number of living cells were seeded exploiting the cell sorting technology (FACS) to exclude dead cells and cellular debris from the initial cell aggregomes (**Fig. 1A**; **Fig. S1E**). The subsequent steps of gastruloid formation assay, including the pulse of CHIR between 48h and 72h AA (**Fig. 1A**), were carried out as previously described (Baillie-Johnson et al., 2015; Turner et al., 2017; van den Brink et al., 2014). Following this protocol, we found that at 96h AA almost 100% of the developing organoids examined displayed an evident protrusion zone, thus indicating a well-preserved timing of gastruloid induction (**Fig. 1B**). Moreover, 24h later (120h AA), two different phenotypes were observed; a prevalent fraction (95-98%) of fully developed elongated-shaped organoids and a minor fraction (2-5%) of elongated organoids but displaying one or more short ectopic protrusions (**Fig. 1C**). Different parameters analysed, including the diameter of aggregomes (48h AA), the length and the volume of the developed organoids (120h AA), displayed a low dispersion around the corresponding mean value (**Fig. 1D**), suggesting a high accuracy of the method. For instance, the diameter of FACS-plated aggregomes fluctuated in a significant lower range (mean=166 μm; min=153 μm; max=180 μm) compared to the handmade aggregomes (mean=156 μm; min=125 μm; max=195 μm) (**Fig. 1D**; **Fig. S1A**). Furthermore, the expression profile of developmental markers that are induced during the aggregome to gastruloid transition in a time-and space-controlled manner (**Fig. 1E**), including *Brachyury* (*T*) (primitive streak, mesoderm), *Sox2* (progenitor cells, stemness), *Sox17* (primitive and definitive endoderm), *Cdx2* (posterior mesoderm), and *Nestin* (neuronal precursor cells), indicated the establishment of cell lineages and antero-posterior (A-P) axis. Moreover, in the attempt to analyse their histological organization, the gastruloids were included in epoxy resin and serially sectioned. Interestingly, the gastruloids generated by naïve TBV2 cells (120h AA) displayed a tightly compacted structure mainly formed by actively dividing cells (**Fig. 1F**), as also confirmed by immunofluorescence analysis for the proliferation marker Ki67 (**Fig. 1G**). Surprisingly, we observed that excessive narrowing of the forward-scattered light (FSC) value of the cells sorted reduced GFE by increasing the fraction of undeveloped aggregomes up to 5-fold (**Fig. S1F**). All together these data pointed to an improved GFE and suggested that a certain degree of phenotypic heterogeneity in the stem cell population is essential to generate functional aggregomes.

**Figure 1.**
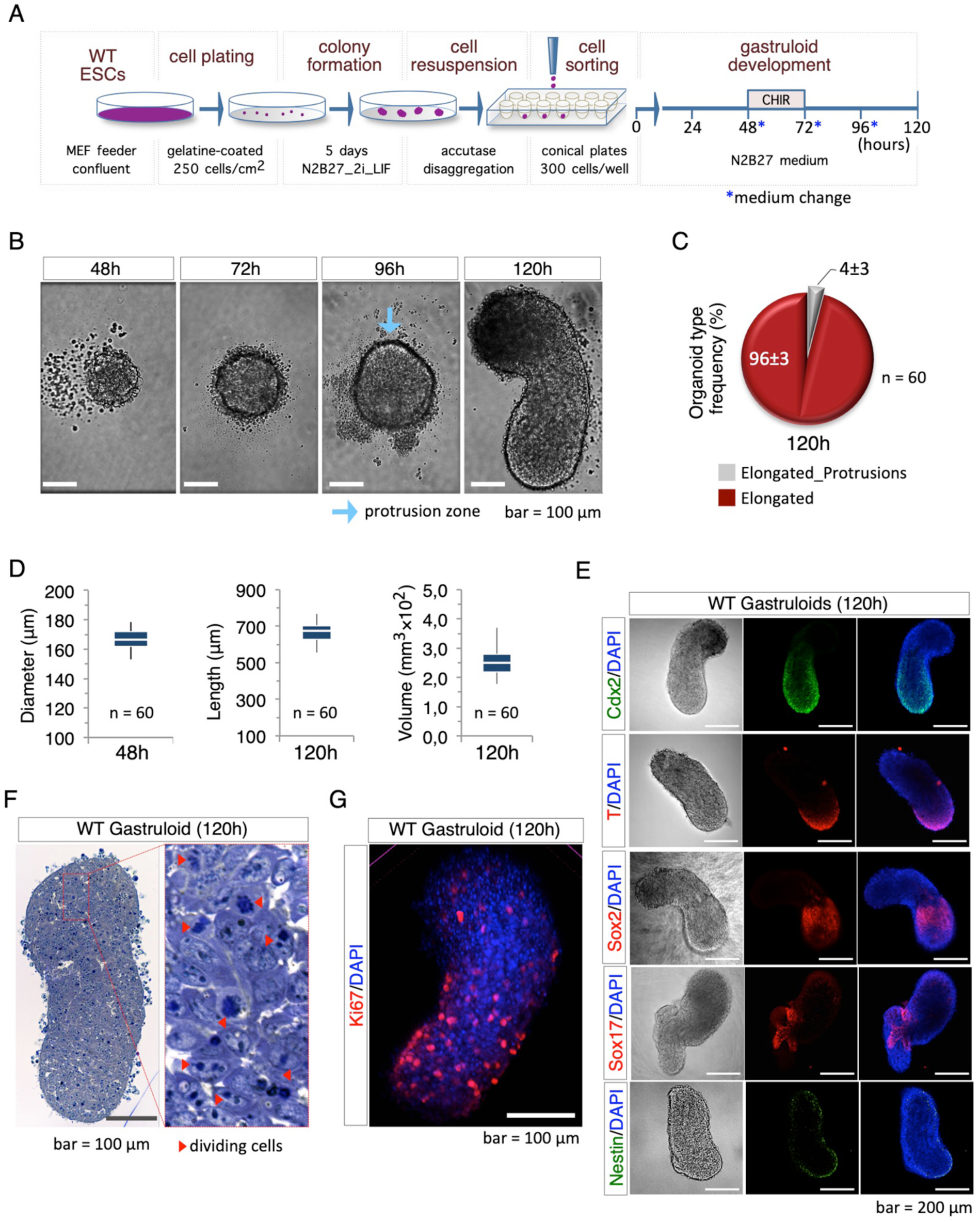
Efficiency of the optimised gastruloid formation assay. (**A**) Schematic representation of the experimental design. Low-density colony formation, accutase dissociation and FACS seeding were introduced to standardize the protocol. 2i_LIF TBV2 mESCs (WT) were seeded in ultra-low attachment 96-well plate and cultured in N2B27 medium for 120h. CHIR was added at 48h. (**B**) Time-course representative bright field images of aggregomes to gastruloids transition showing the morphology of aggregomes from 48h to 120h after aggregation (AA). Light blue arrow indicates the protrusion zone (bar=100 μm). (**C**) Pie chart representing the percentage of the different phenotypic outcomes including organoids without protrusions with a defined AP axis (‘Elongated’) or organoids with protrusions and a defined AP axis (‘Elongated_Protrusions’) (n=60). (**D**) Box plot diagram of the aggregomes diameter at 48h AA (left), and the length (middle) and volume (right) of the gastruloid at 120h AA (n=60/time point). (**E**) Representative pictures of bright-field (left) and confocal immunofluorescence analysis with Cdx2 (green), T/Bra, Sox2, Sox17 (red) and Nestin (green) of 2i_LIF mESC gastruloids at 120h AA. Nuclei were counterstained with DAPI (blue) (bar=200 μm). (**F**) Representative pictures of sections (5-μm thick) of resin embedded gastruloids at 120h AA. The sections were stained with toluidine blue; red arrows show dividing cells (bar =100 μm). (**G**) Representative pictures of confocal immunofluorescence with Ki67 of gastruloid at 120 AA (red). Nuclei were counterstained with DAPI (blue).

We also performed gastruloid formation assays using a feeder-free stem cell line (E14 mESCs). The cells were grown at low density (250 cells/cm^2^) on gelatin-coated plates in naïve-inducing (N2B27/2i/LIF) medium, rising up to 95% of round-domed cell colonies (**Fig. S1G**). The resulting cells were sorted and seeded (300 cells/well) by FACS (**Fig. S1H**), and we have observed that at 120h AA almost all aggregomes became elongated-shaped gastruloids (**Fig. S1I**). Thus, we concluded that our protocol could be applied to both feeder-free and feeder-dependent mESCs.

### *Cripto* is essential for proper gastruloid formation

To validate our experimental approach, we tested the GFE of *Cripto* Knockout (KO) mESCs. *Cripto* (also known as Teratocarcinoma-Derived Growth Factor 1, *Tdgf1*) is a key regulator of pluripotent stem cells (Fiorenzano et al., 2016; Minchiotti, 2005) and is required for the anterior-posterior axis formation during mouse embryo development (D’Andrea et al., 2008; Ding et al., 1998). We compared the GFE of wild-type mESCs (R1) and two independent *Cripto* KO cell lines mESCs (hereafter Cl.#1 and Cl.#2) (Parisi et al., 2003). The cells were seeded at low density on gelatine-coated plates in naive state-inducing medium (N2B27/2i/LIF). Following 5 days in culture, the resulting colonies were dissociated with accutase and the disaggregated cells were seeded in 96- well ultra low attachment plates (300 cells/40 μl) (**Fig. 2A**). At 48h AA both WT and *Cripto* KO cells generated almost spherical cell aggregomes with a diameter of around 160 μm (**Fig. 2B**); the aggregomes of *Cripto* KO Cl.#2 but not Cl.#1 were slightly smaller compared to control (**Fig. 2B**). At 96h AA a protrusion zone became evident in the large majority (~90%) of Control and *Cripto* KO aggregomes (**Fig. S2A**). Later on, at 120h AA, almost all control aggregomes became fully elongated organoids, while a small fraction (<5%) remained as undeveloped spheroids (**Fig. 2C**). Conversely, most of the *Cripto* KO aggregomes (≥95%) maintained an spheroidal morphology (**Fig. 2C**; **Fig. S2A**), providing evidence that in the absence of *Cripto* the aggregome to gastruloid transition was halted. The number of cells required to generate an effective aggregome, *i.e.* able to develop into a fully elongated gastruloid, varies amongst ESC lines (Turner et al., 2017). Thus, to exclude any cell-number effect of *Cripto* KO cells, we used increased number of cells, ranging from 250 to 450 cells/well, to generate the initial aggregomes. Under these conditions, *Cripto* KO aggregomes still failed to elongate (**Fig. S2B**), thus providing evidence that *Cripto* is dispensable for aggregome formation but essential for gastruloid development.

**Figure 2.**
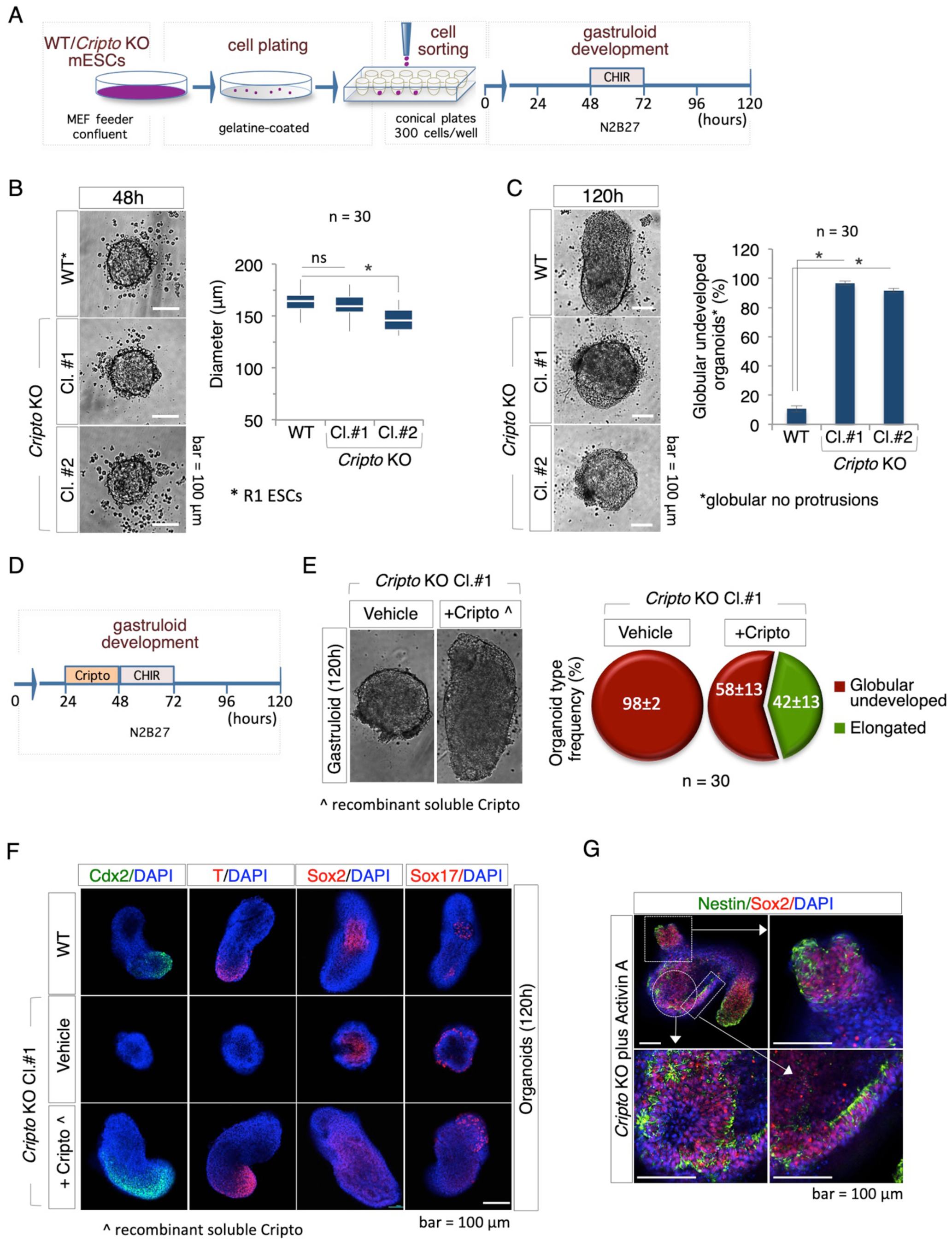
Cripto genetic ablation impairs gastruloids formation. (**A**) Schematic representation of the experimental design. (**B**) Representative pictures (left) and diameter distribution (right) of R1 (WT) and *Cripto* KO mESCs (Cl.#1 and Cl.#2) derived aggregomes at 48h AA (n=30; bar=100 μm). (**C**) Bright field representative pictures (left) of WT and *Cripto* KO ESC-derived gastruloids at 120h AA showing that *Cripto* KO ESCs failed to develop elongated structures. Percentage of undeveloped gastruloids (*globular no protusions) (right) of WT and *Cripto* KO derived organoids at 120h (n=30, *p<0.01, bar=100 μm). (**D**) Schematic representation of the rescue experiment. Soluble Cripto protein (sCripto, 10 μg/ml) was added at 24h AA. (**E**) Bright field representative pictures (left) of *Cripto* KO rescued gastruloids at 120h AA. Pie chart of the percentage of organoid type frequency (right) in *Cripto* KO treated with sCripto or vehicle as Control. (**F**) Representative confocal pictures of immunofluorescence with Cdx2 (green), T (red), Sox2 (red) and Sox17 (red) on from WT and *Cripto* KO ± sCripto -derived organoids at 120h AA. Nuclei were counterstained with DAPI (blue) (bar=100 μm). (**G**) Representative confocal pictures of immunofluorescence with Sox2 (red) and Nestin (green) on *Cripto* KO + ActA derived gastruloids at 120h AA. Nuclei were counterstained with DAPI (blue) (bar=100 μm).

Cripto is a glycosylphosphatidylinositol (GPI)-anchored extracellular protein (Minchiotti et al., 2000), so to further investigate the role of Cripto in gastruloid development we performed rescue experiments by adding a recombinant soluble active form of Cripto (sCripto) at a concentration of 10 μg/ml (Guardiola et al., 2012; Parisi et al., 2003). *Cripto* KO aggregomes were treated with sCripto (from 24 to 48h AA) or left untreated as Control and the GFE was analysed at 120h (**Fig. 2D**). Interestingly, a significant fraction (>40%) of sCripto-treated *Cripto* KO aggregomes developed elongated gastruloids, which appeared even larger in size compared to Control (**Fig. 2E**). We also analysed the expression pattern of different developmental associated genes that are induced during the aggregome to gastruloid transition in a time- and space-controlled manner, including *Brachyury* (*T*), *Cdx2*, *Sox2*, and *Sox17* (Baillie-Johnson et al., 2015; Beccari et al., 2018; Tesar et al., 2007). Immunofluorescence analysis revealed a similar spatial pattern of cells expressing *Brachyury* (*T*) and *Cdx2* in fully elongated WT and *Cripto* KO rescued (Cripto KO + sCripto) gastruloids; which, conversely were almost absent in the *Cripto* KO undeveloped organoids (**Fig. 2F**). The expression domain of both genes in the rescued mutants was larger compared to that of WT gastruloids, suggesting that sCripto treatment expanded the posterior mesoderm fate of gastruloids, which might be due to increased Nodal activity (Minchiotti et al., 2002). Interestingly, *Sox2* and *Sox17* were expressed at different extent in the undeveloped *Cripto* KO aggregomes (**Fig. 2F**; **Fig. S2C**). qPCR analysis performed on 120h AA organoids confirmed and extended these findings e.g *Brachyury* and *Cdx2* were undetectable in *Cripto* KO organoids, whereas their expression was fully rescued in sCripto-treated *Cripto* KO gastruloids (**Fig. S2D**). Expression of the neural marker *Sox2* was significantly higher in *Cripto* KO than in Control (**Fig. S2D**), correlating with the presence of a large group of Sox2 positive cells (**Fig. 2F**) in the central region of the undeveloped *Cripto* KO spheroids. The observation of Nestin positive cells supported the neural fate specification in *Cripto* KO organoids (**Fig. S2C**). All together these findings were in line with previous reports showing that *Cripto* -null mutant embryos mostly consist of anterior neuroectoderm and lack posterior structures (Ding et al., 1998), and provided evidence that *Cripto* is essential for the induction of symmetry breaking (A-P axis formation) in developing mESCs aggregomes.

Cripto is an obligate co-receptor of Nodal (Minchiotti, 2005) and to better define the role of Nodal signaling in the inability of *Cripto* KO cells to generate gastruloids, we investigated the effect of Activin A, a member of the transforming growth factor beta (TGF-β) family of proteins and a potent inducer of Nodal pathway (Pauklin and Vallier, 2015). Activin signaling can be induced in *Cripto* KO mESCs (Fiorenzano et al., 2016). Notably, we found that a transient pulse (24-48h) of Activin A (20 ng/ml) induced the polarization and elongation of a significant fraction (~60%) of *Cripto* KO aggregomes (**Fig. S2E**) which also expressed *Brachyury* at the posterior end of the gastruloid (**Fig. S2F**). However, unlike to what we observed in WT gastruloids, in the elongated gastruloid-like organoids developed by Activin A-treated *Cripto* KO aggregomes, the expression of *Cdx2* and *Sox17* was undetectable (**Fig. S2F**), whereas large zones of *Nestin* (neuronal precursor marker) expressing cells were observed, thus indicating a deregulation of neural differentiation (**Fig. 2G**; **Fig. S2F**). These data suggest that Activin A rescues WT polarization and elongation but fails to induce proper gastruloid development in *Cripto* KO aggregomes. Moreover, most of the organoids generated by *Cripto* KO cells treated with Activin A displayed ectopic-protrusions. This might likely reflected the size of the Activin A-treated aggregomes (48h AA), which was consistently larger (>200 μm) compared to the aggregomes that elongate properly. Thus, the abortive development of *Cripto* KO aggregomes validated our experimental approach and further supported the idea that gastruloid development mimics early embryo gastrulation.

### Primed pluripotency impairs gastruloid development

In order to investigate the impact of pluripotency on GFE, we first set up the *in vitro* conditions useful to capture mESCs into different primed states of pluripotency. We thus induced the naive to primed transition by treating mESCs with Fibroblast Growth Factor (12 ng/ml) and Activin A (20 ng/ml) (F/A mix), after LIF withdrawal (Ying et al., 2008). As schematized in **Fig. 3A**, naïve (2i_LIF) TBV2 ESCs were seeded at low density (1500 cells/cm^2^) on serum-coated plates and incubated in N2B27/1% Knockout Serum Replacement (KSR) medium supplemented with F/A. F/A-induced cells were cultured for two (2d_F/A) and five (5d_F/A) days and the resulting cells were characterized at molecular and phenotypic level. First, we analyzed the expression profile of different pluripotency and lineage specific markers by qPCR. As expected, the pluripotency markers *Nanog*, *Sox2*, *Rex1* and *Dppa5a* were expressed at lower levels in 2d_F/A cells compared to naïve ESCs, whereas the differentiation markers *Apelin* receptor (*Aplnr* or *ApJ*), *Cerberus* (*Cer1*), and *Brachyury* (*T*) were overexpressed (**Fig. 3B**). Conversely, 2d_F/A cells expressed higher levels of pluripotency and lower levels of differentiation markers compared to 5d_F/A cells. To further characterize 2d_F/A and 5d_F/A cells, we assessed their ability to revert to the naïve state by performing colony formation assays. To this end, F/A-treated cells were dissociated with accutase and plated at low density on gelatin-coated plates in naïve-inducing (N2B27/2i/LIF) medium. Under such conditions, 2d_F/A but not 5d_F/A cells, grew generating naïve-like domed-shaped colonies (**Fig. 3C**; **Fig. S3A**). A key functional feature that distinguishes early-primed Epiblast-like cells (EpiLC) from primed Epiblast Stem Cells (EpiSCs) and naïve ESCs is the ability to generate primordial germ cells (PGCs). We thus assessed the PGC differentiation potential of F/A-treated cells applying a previously described protocol (Hayashi et al., 2018). As shown below (see **Fig. 5**), 2d_F/A but not 5d_F/A cells, efficiently differentiate into PGCs. Together these findings suggest that 2d_F/A cells have acquired features of early-primed EpiLCs, including spontaneous reversion to naïve state and differentiation into PGC; whereas, 5d_F/A cells exhibited features of primed EpiSCs, including failure to revert to naïve state and inefficient PGC differentiation (Morgani et al., 2017).

**Figure 3.**
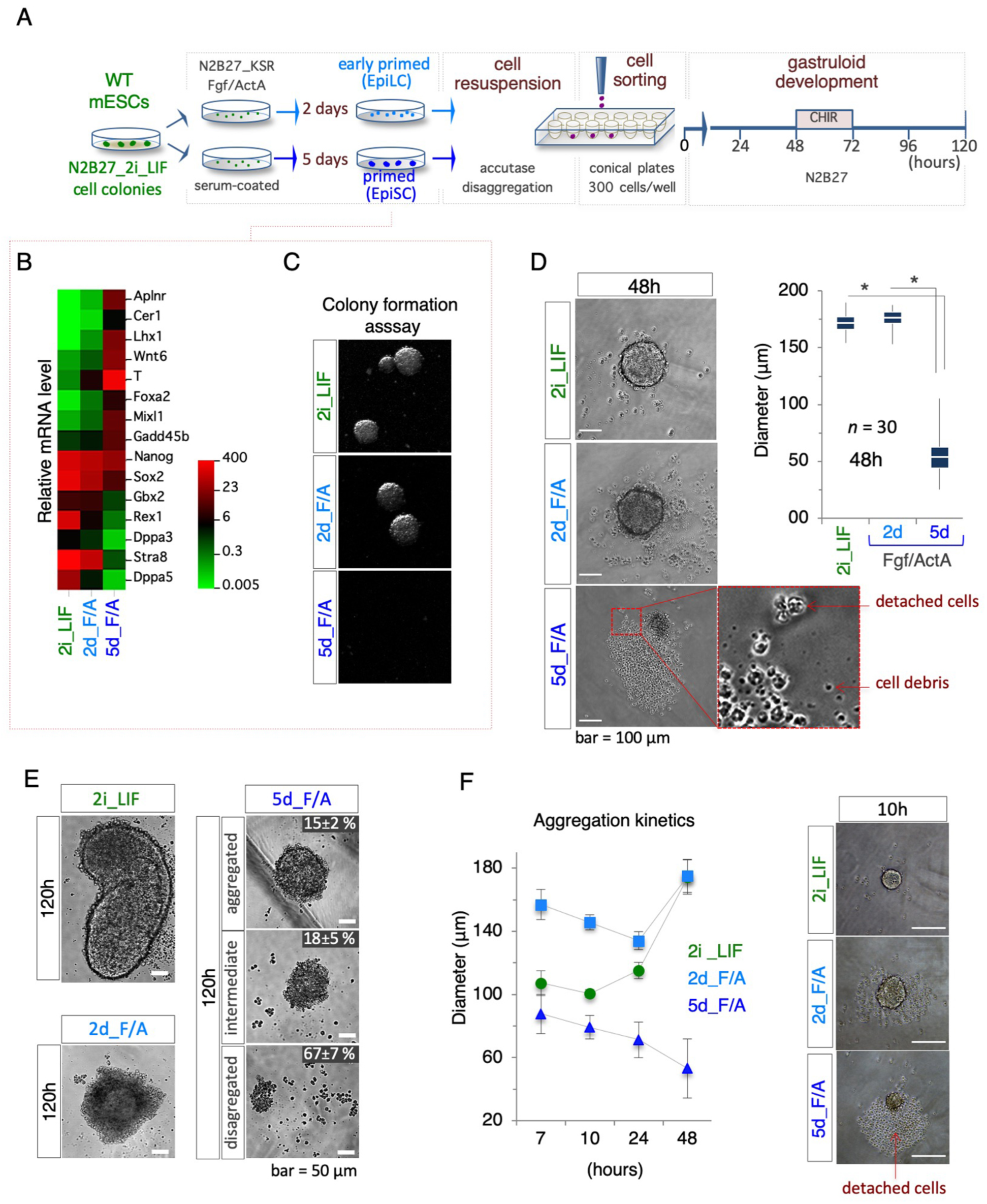
F/A primed pluripotent cells fails to generate gastruloids. (**A**) Schematic representation of the experimental design using EpiLCs (2d_F/A) and EpiSCs (5d_F/A) cells. (**B**) Heat-map of mRNA levels of the indicated genes in naïve (2i_LIF), EpiLCs and EpiSCs cells. (**C**) Colony formation assay of naïve, EpiLCs and EpiSCs cells. (**D**) Bright field representative pictures of different aggregomes at 48h AA (left) and box plot diagram (right) of diameter distribution. Red arrows indicate detached cells and debris (n=30; bar=100 μm). (**E**) Representative bright field images of (left) 120h AA organoids and (right) different aggregomes phenotypes with percentage at the indicated conditions (left). (**F**) Aggregation kinetics (left) of naïve (2i_LIF), EpiLCs (2d_F/A) and EpiSCs (5d_F/A) aggregomes measured by diameter distribution (μm). Representative bright field images of aggregomes at 10h generated from the indicated cells (right). Red arrow shows detached cells.

Once the pluripotency state of 2i_LIF (naïve, Control), 2d_F/A (early-primed EpiLCs) and 5d_F/A (primed EpiSCs) has been ascertained, we used the optimized gastruloid formation protocol to compare their GFE (**Fig. 3A**). First, we found that naïve and primed cells showed remarkable differences in both the cytometric parameters (**Fig. S3B**), and the ability to generate aggregomes (**Fig. 3D**). Indeed, at 48h AA, both 2d_F/A (EpiLCs) and 2i_LIF (naïve) cells generated aggregomes similar in shape (spheroidal) (**Fig. 3D**, left) and size (diameter mean ranging around 170-180 μm) (**Fig. 3D**, right). In contrast, the aggregomes generated by 5d_F/A (EpiSCs) were significantly smaller in size (~50 μm) (**Fig. 3D**). Later on, at 120h AA, while Control aggregomes developed fully elongated gastruloids (**Fig. 3E**; **Fig. S3C**), quite unexpectedly, almost all the aggregomes generated by EpiLCs maintained an spheroidal morphology, with a central globular aggregate of cells surrounded by a disorganized and irregular tissue-like structure (**Fig. 3E**; **Fig. S3C**), suggesting that they failed to undergo proper aggregome to gastruloid transition. Finally, 5d_F/A (EpiSCs) aggregomes, which were extremely small at 48h AA, became only abortive structures whose morphology ranged from spheroidal cell aggregates (~15%) to skimpy irregular cell clusters (~70%) (**Fig. 3E**). Since none of the EpiLC- and EpiSC-derived aggregomes scored (n ≥300) was able to generate an elongated gastruloid-like organoid, we concluded that 2 days incubation in primed state-inducing medium (N2B27_KSR plus F/A mix and without LIF), were sufficient to abolish the gastruloid formation ability of naïve mESCs. Surprisingly, after transient LIF supplementation (from 0 to 48h AA) more than 70% of the EpiLCs aggregomes generated a gastruloid-like organoids at 120h AA (**Fig. S3D-F**). All together these data indicated that, under our experimental conditions, primed stem cells were unable to generate efficient aggregomes (GFE value = 0%).

### Primed pluripotency state reduces cell aggregation propensity

The precise dimension of the mESCs aggregomes (48h AA) is a critical parameter for gastruloid induction (Baillie-Johnson et al., 2015); thus the abortive development of the small aggregomes generated by 5d_F/A (EpiSCs) was almost expected. Reduced aggregome size can be a consequence of either a defective cell aggregation process or a disaggregation subsequent to a normal aggregation process. To investigate this issue directly, we imaged aggregome generation at early time points. Recent findings reported that under low-attachment conditions mESCs (250-300 cells/well) spontaneously aggregates in 8-10h (Turner et al., 2017); in line with this observation, we found that already 7h after seeding 2i_LIF (naïve) TBV2 cells had generated irregular-shaped cell aggregates (**Fig. S3G**). Later on (10h) we observed that the aggregates of naïve cells became highly compacted and displayed a spheroidal-shaped morphology (mean diameter ~100 μm) (**Fig. 3F**). Within this time window only few free non-adherent cells were observed, thus confirming the extraordinary propensity of naïve cells to aggregate and generate stable cell-cell interactions. Similarly, early aggregomes (7h after seeding) of d2_F/A (EpiLCs) were spherical, although less compacted (mean diameter ~150 μm) and surrounded by a few number of non-adherent cells (**Fig. 3F**). By 10h onwards, the dimension of EpiLC (d2_F/A) aggregomes decreased, reaching its minimal (~135 μm) around 24h after seeding and without increasing the number of detached cells (**Fig. 3F**). It was thus reasonable to speculate that the generation of gastruloid-developing aggregomes requires the compaction/condensation, most likely through the stabilization of cell-cell contacts, of the initial cell aggregates. Conversely, 7h after seeding 5d_F/A (EpiSCs) aggregates were much smaller in size (50-70 μm) and, most relevant, were surrounded by a large number of non-adherent cells and cellular debris (**Fig. 3F**; **Fig. S3G**). We also analysed a previously reported gene expression profile of 5d_F/A treated TBV2 cells (GEO accession: GSE84373) and found that their reduced aggregation ability correlates with a significant up-regulation of focal adhesion-related genes (**Fig. S3H**). These data suggested that during the ESC to EpiSC transition TBV2 cells undergo a substantial loss of their capacity to generate stable cell-cell contacts, and this likely resulted in failure to generate aggregomes.

### Proline-induced cells undergo premature gastruloid development

To further evaluate the impact of primed pluripotency on gastruloid development, we assayed the GFE of L-Proline-induced cells (PiCs) (Casalino et al., 2011; Comes et al., 2013), also known as early primitive ectoderm-like cells (EPL) (Washington et al., 2010). PiCs were obtained by seeding naïve (2i_LIF) mESCs at low clonal density on gelatin-coated plates in DMEM/FBS medium supplemented with LIF and L-Proline (500 μM), as previously described (Comes et al., 2013). As expected, more than 90% of the cell colonies displayed a typical flat and irregular morphology after 4-5 days in culture (**Fig. S4A**). PiCs were thus dissociated by accutase and the resulting cells were seeded by FACS and incubated to induce gastruloid formation (**Fig. 4A**). Indeed, untreated Control and PiCs cells displayed different flow cytometric features (FSC and SSC), indicating that a high L-Proline regimen modified ESCs size and granularity (**Fig. S4B**). Two-day-old aggregomes (48h AA) generated by PiCs were smaller in size (mean diameter = 120 μm) compared to naïve (2i_LIF) ESCs aggregomes (mean diameter = 160 μm) (**Fig. 4B**). Of note, these differences persisted also after increasing the number of seeded cells from 250 up to 350 (**Fig. S4C**). Remarkably, by 72h onwards, PiCs aggregomes started to undergo elongation; whereas, as expected, control aggregomes still exhibited a round-shaped morphology (**Fig. 4C**). Thus, elongation of PiCs aggregomes’ was anticipated by 24-36 hours (72-96h AA PiCs vs 96-120h AA untreated ESCs); yet the overall dimension of the gastruloids-like organoids generated by PiCs was smaller compared to control cells (**Fig. 4C**). These findings suggested that PiCs aggregomes were much more prone to undergo symmetry breaking and elongation than normal naïve-derived aggregomes. Later on, at 96h AA, two different morphologies of PiCs-derived organoids were observed; indeed, the majority (~60%) displayed a gastruloid-like fully elongated shape, while the remaining (~40%) maintained the spheroidal morphology of the initial aggregomes (**Fig. 4D**). The mean length of PiCs-derived gastruloids was up to 2 times lower compared to control gastruloids (**Fig. S4D**).

**Figure 4.**
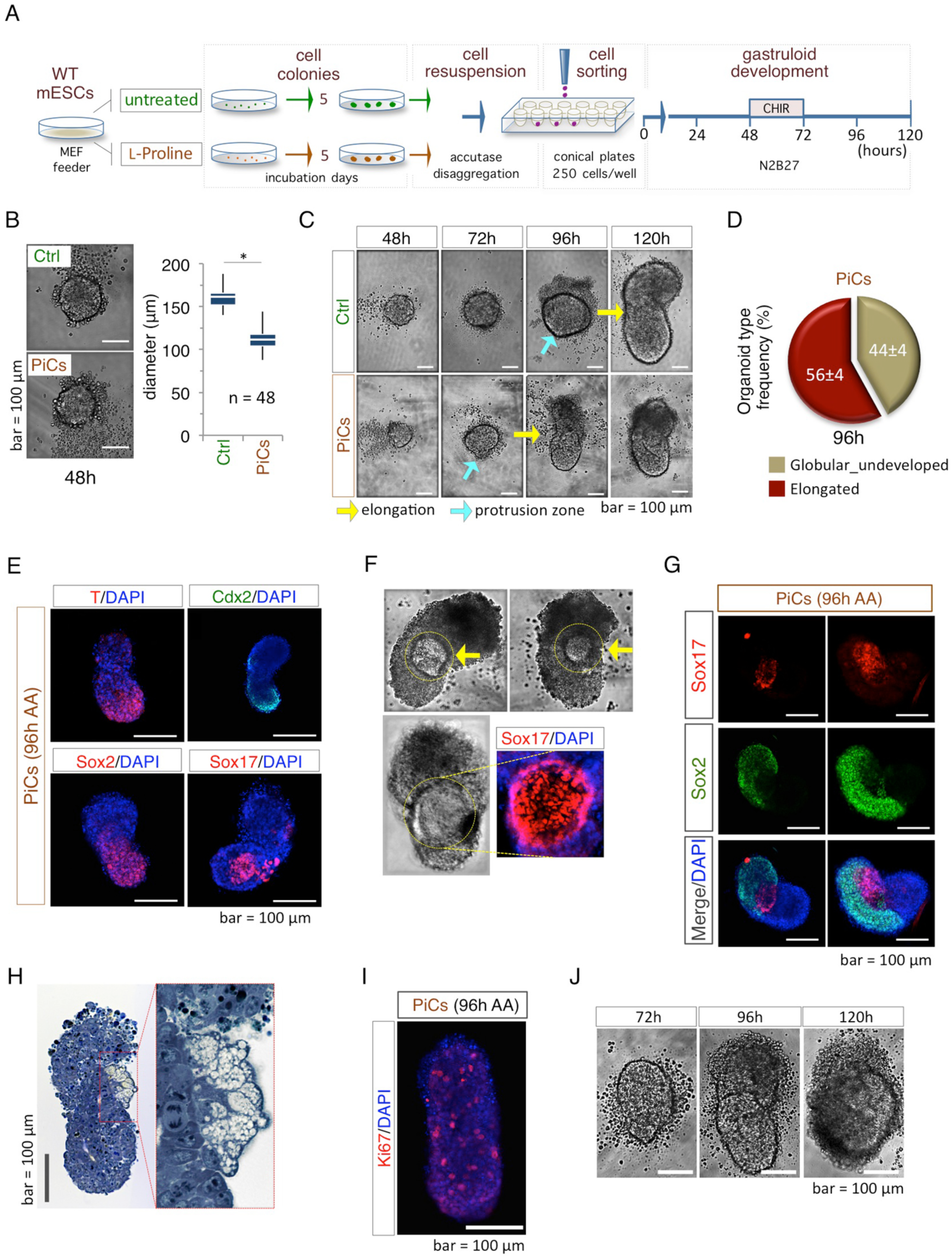
L-Proline-treated ESCs (PiCs) are competent for gastruloids formation. (**A**) Schematic representation of experimental scheme using L-Proline-treated cells (PiCs) and untreated control (Ctrl). (**B**) Bright field representative pictures (left) of Ctrl and PiCs aggregomes at 48h AA and box plot diagram (right) of diameter distribution (n=48; bar=100 μm). (**C**) Bright filed representative pictures of the aggregome to gastruloid transition of PiCs and Ctrl cells at the indicated time points (bar=100 μm). Light blue and yellow arrows indicate the protrusion zone, and the switch from ovoidal to elongated shape, respectively. (**D**) Pie chart of PiCs derived organoids’ types frequency. (**E**) Representative confocal pictures of immunofluorescence with T/Bra (red), Cdx2 (green), Sox2 and Sox17 (red) on PiCs derived gastruloids at 96h AA. Nuclei were counterstained with DAPI (blue) (bar=200 μm). (**F**) Bright-field representative pictures of different PiCs derived gastruloids at 96h showing the Sox17 positive area (yellow circle). (**G**) Representative confocal pictures of immunofluorescence with Sox17 (red) and Sox2 (green) on PiCs derived gastruloids at 96h AA. Nuclei were counterstained with DAPI (blue) (bar=100 μm). (**H**) Representative pictures of blue toluidine staining on sections of PiCs derived gastruloids at 96h AA. Enlargement shows a differentiated area (bar=100 μm). (**I**) Representative pictures of confocal immunofluorescence with Ki67 of PiCs derived gastruloid at 96h AA (red). Nuclei were counterstained with DAPI (blue) (bar=100 μm). (**J**) Bright field representative pictures of PiCs-derived gastruloids at the indicated time points (bar=100 μm).

To evaluate if this peculiar developmental process (advanced lengthening and reduced dimension) correlated with altered expression of key developmental genes, we analysed the expression pattern of different markers by IF analysis. *Cdx2* and *Brachyury* were both expressed in specific territories of PiCs elongated organoids (**Fig. 4E**). Frequently, the PiCs-derived organoids exhibited a prominent round-shaped zone of histological heterogeneity, which was usually located in the middle/central region of the gastruloid, and stained positive for the mesoendodermal marker *Sox17* (**Fig. 4F**). Interestingly, large areas of cells expressing the pan-neuronal marker *Sox2* were observed adjacent to and surrounded the central region of *Sox17* expressing cells (**Fig. 4G**). To better analyze their histological organization, resin-embedded PiCs-derived gastruloids (96h AA) were sectioned and stained with toluidine blue. Histological examination revealed the presence of both actively dividing cells, as expected in a developing gastruloid, but also areas of differentiated cells/tissues (**Fig. 4H**). Immunofluorescence analysis of Ki67 confirmed the presence of actively dividing cells (**Fig. 4I**). At 120h AA a significant fraction (>50%) of PiCs-derived organoids showed a disorganized structure surrounded by a large number of detached cells (**Fig. 4J**; **S4E**), suggesting that underwent early disaggregation compared to naïve-derived control gastruloids. All together these findings support the idea that L-Pro supplementation in the presence of LIF forced the cells towards an early and reversible primed state of pluripotency, which maintained the competence to generate gastruloid-like organoids.

### PiCs are competent to PGC-like cells differentiation

Our quite unexpected findings that early-primed EpiLCs (2d_F/A) were able to generate properly shaped but unproductive aggregomes; *i.e.* unable to undergo aggregome to gastruloid transition; whereas, PiCs (L-Proline/LIF) were able to generate small but competent aggregomes (**Fig. 3**), prompted us to further investigate the functional features of EpiLCs and PiCs. It is known that the ability to differentiate into cells displaying features of primordial germ cells, named primordial germ cell-like cells (PGCLCs), is a unique feature of EpiLC cells, whereas both naïve and EpiSCs cells are recalcitrant to undergo PGC differentiation (Morgani et al., 2017). We thus investigated if PiCs were competent to generate PGCLC compared to naïve and primed (EpiLCs and EpiSCs) pluripotent cells as control. Naïve (2i/LIF) cells were used to generate early-primed (2d_F/A, EpiLC), primed (5d_F/A, EpiSC), and PiCs (L-Proline/LIF) as schematized in **Figure 5A**. The cells were dissociated with accutase and seeded (2500 cells/well) in ultra-low attachment 96-well conical plates to force aggregation (**Fig. 5A**). The resulting aggregomes were incubated in GK15 medium supplemented with a standard mix of PGC-inducing growth factors, including Bone Morphogenetic Protein 4 (BMP4), Epidermal Growth Factor (EGF), Stem Cell Factor (SCF), and LIF, as previously described (Hayashi et al., 2013). After 4 days of incubation it was observed that the size of aggregomes vary among the different pluripotent states analysed (**Fig. 5B**). Indeed, the aggregomes generated by PiCs were smaller compared to those generated by naïve and early-primed EpiLCs (**Fig. 5B**). Most relevant, as already observed performing the gastruloid formation assay (**Fig. 3**), 5d_F/A (EpiSCs) were refractory to generate cell aggregates (data not shown). PGCLC are identified/marked by the co-expression of *Blimp1* and *Oct4* genes (Hayashi et al., 2018), and immunofluorescence analysis revealed the presence of *Blimp1*/*Oct4* double positive cells in aggregomes generated by EpiLCs and PiCs (**Fig. 5C**). To corroborate these results, we analysed the expression profile of different markers including *Prdm14* and *Nanos3* by q-PCR assays. Remarkably, all the genes analysed were significantly induced in EpiLCs- and PiCs-derived aggregomes, including *Blimp1* (**Fig. 5D**). Moreover, EpiLCs- and PiCs-derived aggregomes showed a high percentage of AP2-γ (Transcription factor AP2-γ encoded by *Tfap2C* gene) positive cells compare to 2i_LIF-derived aggegomes. All together these findings indicate that both EpiLCs and PiCs are competent to PGCLC differentiation, and provide further evidence that a high L-Proline regimen induce an early-primed state of pluripotency, which exhibits the unique competence for both gastruloid formation and differentiation into PGC-like cells.

**Figure 5.**
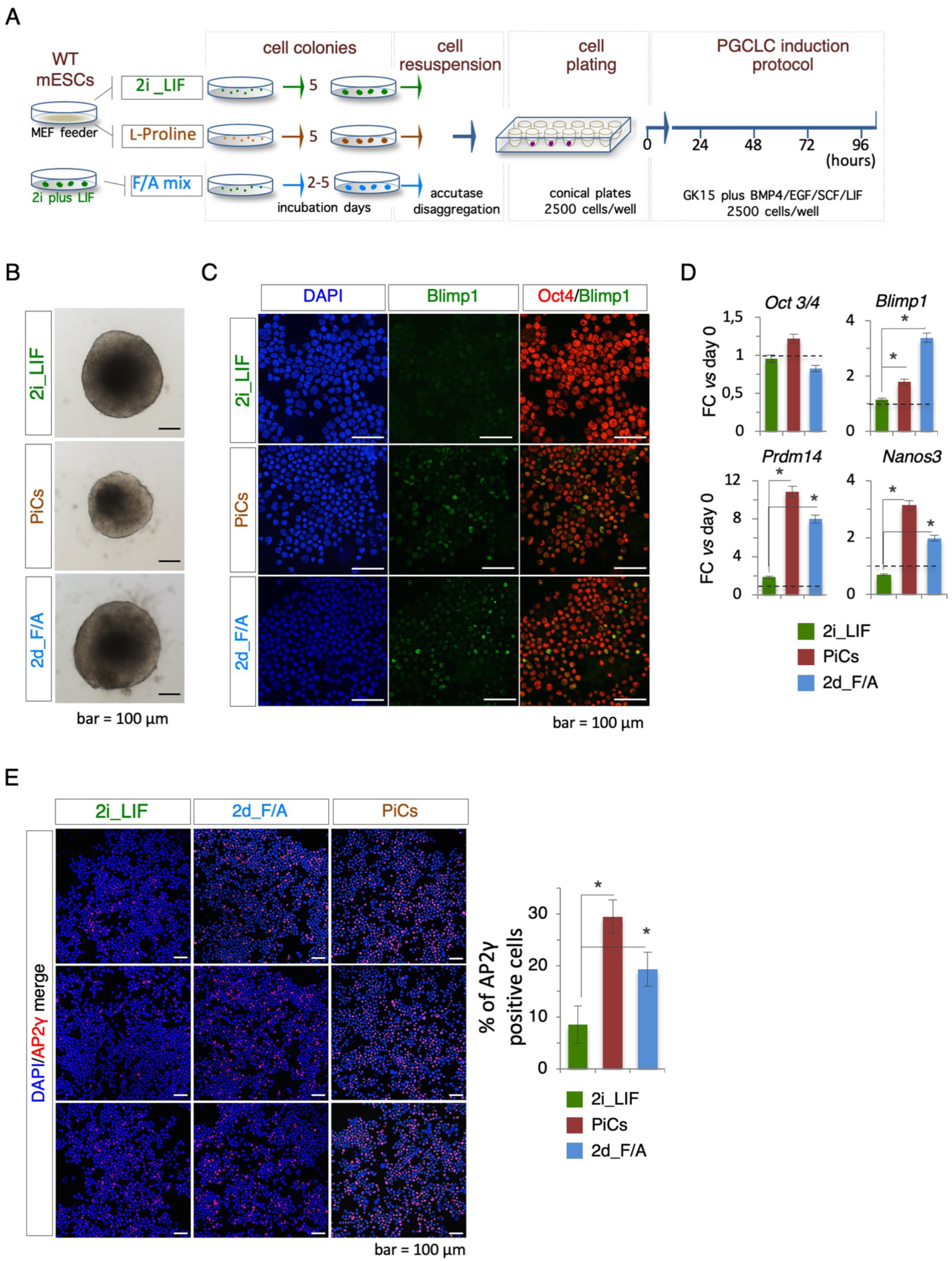
Primordial germ cell-like cells (PGCLCs) competence induction discriminate different pluripotent states. (**A**) Schematic representation of the experimental strategy to induce PCGLCs differentiation using naïve (2i_LIF), EpiLCs (2d_F/A) and EpiSCs (5d_F/A) cells to perform the PCGLCs induction. (**B**) Bright field representative pictures of 96h cell aggregomes derived from naïve, PiCs, and EpiLCs cells, generated as indicated in panel (A) (bar=100 μm). (**C**) Representative confocal pictures of immunofluorescence with Oct4 (red) and Blimp1 (green) on cytospined cells derived from aggregomes of naïve, PiCs and EpiLCs. (**D**) RT-qPCR analysis of *Oct3/4*, *Blimp1*, *Prdm14* and *Nanos3* genes in the indicated conditions. Data are normalized to *Gapdh* and are mean ± SD and represent of fold change *vs* 0h (n=3; p<0.001). (**E**) Representative confocal pictures of immunofluorescence with AP2γ (red) (left) and percentage (%) of AP2γ positive cells at indicated conditions (right) (n=3; p<0.01).

## DISCUSSION

Stem cell pluripotency can be examined at a molecular and/or phenotypic level. However, while in recent year the development of global molecular profilings’ applications (i.e. transcriptomics, epigenomics, metabolomics, proteomics) have provided comprehensive sets of pluripotency state-associated molecular signatures (Habibi et al., 2013; Pijuan-Sala et al., 2019), the functional assays used to define the different pluripotency state-associated phenotypes remained unchanged and currently require the use of *in vivo* models (blastocyst colonization and teratome formation competence). For instance, the contribution to chimeras is considered the gold-standard method to discriminate naïve (ground state) from primed mESCs (Mascetti and Pedersen, 2016), and results can be biased by the heterogeneity of the population analysed and the number of cells injected into blastocysts. Similarly, in the teratoma formation assay the identification of cells derived from the three primary germ layers into histological complex tumours is imprecise. In this context, an alternative emerges from the gastruloid formation efficiency (GFE) assay proposed here (**Fig. 1**; **Fig. S1**). Certainly, GFE is a cost-effective, quantitative, animal-free and robust experimental approach to distinguish unambiguously ESCs in naïve (GFE>95%) and primed (GFE=0) state of pluripotency.

For a genetic validation of the system we have used *Cripto* KO mESCs, revealing that *Cripto* ablation impairs the induction of A-P axis development, and thus the aggregome to gastruloid transition (**Fig. 2**). Our results are in line with previous reports showing that *Cripto*-null mutant embryos die at d7.5 due to their inability to gastrulate (Ding et al., 1998). Of relevance, the expression of the endoderm marker *Sox17* in undeveloped *Cripto* KO organoids (**Fig. 2**), supports the idea that *Cripto* is dispensable for endodermal fate induction (Jin and Ding, 2013). Moreover, our observation that Cripto and Activin A proteins are both able to re-establish WT polarization and elongation in *Cripto* KO aggregomes (**Fig. 2**; **Fig. S2**), correlates with the observation that both these proteins restore the WT phenotype of zebrafish *oep* (one-eyed pinhead) mutants (Gritsman et al., 1999). Therefore, based on the following considerations: i) Cripto makes mESCs competent to respond to Nodal (Parisi et al. 2003), ii) Nodal is essential for gastruloid development (Turner et al., 2017) and, iii) Activin A can activate the same receptors and effectors of Nodal (Pauklin and Vallier, 2015), it is reasonable to speculate that a Cripto-Activin/Nodal pathway underlies the induction of A-P axis formation during gastruloid development.

Our results highlight a further concern regarding the role of *Cripto* in gastruloid development. Certainly, unlike the WT gastruloids, the elongated organoids generated by the Activin A-treated *Cripto* KO aggregomes are made mostly of cells expressing neural markers (*Sox2* and *Nestin*) (**Fig. 2**). This correlates with the observation that genetic ablation of *Cripto* results in increased neuralization and dopaminergic differentiation of mESCs (Lonardo et al., 2010; Parisi et al., 2003). Thus, our data indicates that Activin A overcome only in part the lack of *Cripto* in gastruloids development, and suggest that Cripto, in an Activin A-unrelated manner, is required to restrain the formation of neural precursors during gastruloid development.

We have exploited the potential of the improved gastruloid formation assay to evaluate the impact of the pluripotency state on GFE, and demonstrate that unlike 2i_LIF (naïve) cells, dissociated primed 5d_F/A (EpiSCs) fail to re-aggregate and thus to generate proper aggregomes of correct size and shape (**Fig. 3**). In line with these findings, Hayashi and co-workers recently showed that F/A treatment strongly weakens the intrinsic propensity of naïve mESCs to aggregate, which eventually leads to failure of differentiation in Primordial Germ Cell-like Cells (PGCLCs) (Hayashi et al., 2011). Indeed, already after 3 days of F/A treatment only small and poorly compacted cell aggregates were generated (Hayashi et al., 2011). Notably, re-aggregation of dissociated/singularized cells, *i.e.* the formation of appropriate aggregomes, is a critical initial event for development of gastruloid (Baillie-Johnson et al., 2015; Turner et al., 2016; Turner et al., 2017) and many different kinds of organoids (Takebe et al., 2015; Takebe et al., 2012). The molecular mechanism(s) underlying aggregome formation, and the reason for a reduced re-aggregation competence of EpiSCs remains unknown. It is known that cell adhesive interactions rely on the expression and affinities of adhesion molecules, and we showed that a significant up-regulation of focal adhesion-related genes occurs in TBV2 ESCs after 5 days of F/A treatment (EpiSCs) (**Fig. S3**). In correlation, primed TBV2 ESCs (5d_F/A) preferentially generate flat-instead of domed-shaped cell colonies, thus suggesting a high propensity to generate cell-substrate rather than cell-cell adhesive interactions. Moreover, E-Cadherin (encoded by *Cdh1*) plays an essential role in cell-cell adhesion, and it has been reported that forced expression of *Cdh1* improved the chimeric formation ability of EpiSCs (integration into the ICM after blastocyst injection) (Ohtsuka et al., 2012). Thus, it is reasonable to speculate that the loss of re-aggregation ability of EpiSCs is caused by an F/A-induced down-regulation, activity inhibition and/or delocalization of cell-cell adhesion molecules such as E-Cadherin.

After 2 days of F/A treatment in the absence of LIF and 2i mix (early primed EpiLCs), the cells maintain the capacity to aggregate, even though the aggregomes generated fail to undergo morphological elongation and gastruloid development (**Fig. 3**). The reason of developmental failure of EpiLCs aggregomes remains unknown. Compaction/stiffening is required for the intrinsic induction of TGFβ signalling and chondrogenic differentiation in mesenchymal stem cell aggregates (Sarem et al., 2019), and we have observed that aggregomes of EpiLCs display a less compacted structure compared to naïve-derived aggregates (**Fig. 3**). Thus, it is tempting to reason that reduced compaction efficiency might generate abortive aggregomes by preventing the intrinsic activation of a crucial signalling pathway(s). Turner and co-workers showed that gastruloid development strictly relies on the activation of WNT and Nodal signalling pathways, whereas BMP pathway appears dispensable (Turner et al., 2017). Since a WNT signalling agonist is exogenously provided during GFE assay, it is possible that alteration of Nodal signalling activation could underlie, at least in part, the EpiLC phenotype. In line with this idea, EpiLCs (**Fig. 3**), *Nodal* KO cells (Turner et al., 2017) and *Cripto* KO cells (**Fig. 2**) generate aggregomes displaying a similar undeveloped phenotype.

L-Proline-induced cells (PiCs) exhibit some features of the early-primed EpiLCs. Specifically, PiCs generate teratomas and colonize mouse blastocyts and, spontaneously and efficiently revert to naïve pluripotency state (Casalino et al., 2011; Comes et al., 2013; D’Aniello et al., 2017b), and here we have unveiled that PiCs are able to differentiate into PGCLCs as well as EpiLCs (**Fig. 5**). However, while EpiLCs are in a non-culturable/transient pluripotency state, PiCs are stably captured in a LIF-dependent early-primed state of pluripotency, and we found that EpiLCs generate only abortive aggregomes (**Fig. 3**), whereas PiCs generate effective aggregomes (**Fig. 4**). We ascribed the developmental failure of EpiLCs with a loss of pluripotency. Certainly, the early-primed pluripotency state of EpiLCs is achieved incubating mESCs in the absence of LIF and 2i mix, whereas early-primed pluripotency state of PiCs is captured incubating mESCs with LIF. A key role of pluripotency in GFE is supported by the observation that LIF supplementation during the first 2 days of incubation, *i.e.* before A-P axis induction with WNT signalling agonist CHIR, is sufficient to restore normal elongation in EpiLC aggregomes (**Fig. S3**).

PiCs aggregomes elongate earlier than control, generating elongate-shaped gastruloid-like organoids, which express markers of A-P axis formation (**Fig. 4**). Why PiCs aggregomes undergo symmetry breaking before control is unknown. However, it has been shown that cultured PiCs are heterogeneous with a large majority of cells displaying features of an intermediate state of pluripotency, and two minor fractions of cells displaying, respectively, features of the extreme naïve or primed states (D’Aniello et al., 2017b). Thus, it is reasonable to speculate that the aggregation of cells at different state of pluripotency may contribute to anticipate the aggregome to gastruloid transition.

In conclusion, here we report an optimized *in vitro* gastruloid formation efficiency (GFE) assay, useful to functionally characterize mouse stem cells pluripotency. Besides being a valid (low-cost, animal-free) alternative to the *in vivo* chimera assay, the particular configuration (FACS combined with, 96 well-plates) and robustness (output >95%) of our GFE assay makes it suitable for large-scale (high-throughput) phenotype-based genetic screenings (shRNA, miRNA) for the identification of genes involved in gastruloids/embryo development. Moreover, the screening of metabolites and drugs libraries could allow the identification of beneficial/toxic metabolites and potential teratogens.

## MATERIALS AND METHODS

### Culture of ESCs, reagents, and treatments

Wild type mouse TBV2 (129/SvP) and R1 ESCs and two independent *Cripto* KO ESC clones, DE7 and DE39 (named Cl.1 and Cl.2) (Parisi et al., 2003; Persico et al., 2001), were used throughout the study. ESC lines were maintained on a feeder layer of mitomycin C-treated primary MEFs according to standard procedures. Feeder-independent ESCs were cultured onto gelatin-coated plates. ESCs were cultured in high glucose Dulbecco’s modified Eagle’s medium (DMEM, Invitrogen, Life Technologies) supplemented with 15% ES-screened Fetal Bovine Serum (FBS, Euroclone), 0.1 mM β-mercaptoethanol (Sigma-Aldrich), 1 mM sodium pyruvate, 2 mM glutamine, 100 U/ml penicillin/streptomycin (all from GIBCO), and 1000 U/ml recombinant LIF (ESGRO, Millipore). 2i Medium (N2B27) was supplemented with PD0325901 (1 μM), CHIR99021 (3 μM) and LIF (ESGRO, Millipore). Recombinant soluble Cripto protein (sCripto) was used at the concentration of 10μg/ml. All cell lines were routinely tested and confirmed to be free of mycoplasma.

### *In vitro* capture of primed state

For ESC to EpiLC and EpiSC transition, 2i mESCs were plated at 1500 cells/cm^2^ onto FBS-coated plates and grown in N2B27 medium supplemented with 20 ng/ml Activin A (Invitrogen) and 12 ng/ml bFgf (Provitro) and cultured for 2 days (2d_F/A, EpiLCs) and 5 days (5d_F/A, EpiSCs), respectively.

### *In vitro* generation of L-Proline-induced cells (PiCs)

To induce ESC to PiC transition ESCs were plated at low density (50–250 cells/cm^2^) in complete medium (DMEM/15%FBS/LIF) on gelatin-coated plates and grown for 5 days in the presence/absence of L-Pro (250–500 μM) (Sigma-Aldrich), with a medium change at day 3. PiCs were harvested using accutase (Sigma-Aldrich).

### Colony type quantification

ESC colonies were washed twice with phosphate-buffered saline (PBS) and fixed/stained with a solution of PBS 1x/6% glutaraldehyde/0.15% crystal violet. After 30-40 min at RT, the stained cell colonies were carefully washed with tap water and dried for further analysis. The colony type percentage (domed/flat) was determined by quantifying the number of domed and flat colonies over the total number of colonies. This analysis was performed blinded by two investigators on at least 400 colonies per condition.

### Gastruloid formation assay

For gastruloids formation assay we performed an optimised version of the previously published protocol (Baillie-Johnson et al., 2015; van den Brink et al., 2014). Briefly, cells grown at low density, the resulting colony were dissociated with accutase (Sigma-Aldrich) and then an exact number of cells (2.5×10^2^ or 3.0×10^2^) were FACS sorted/seeded in U-shaped ultra-low attachment 96-multiwell (Corning Costar) and allowed to aggregate for 48h in N2B27 medium without supplemental growth factors. After 48h in culture, CHIR was added at the concentration of 3μM and maintained for 24 hours. From 72h onwards, N2B27 medium (150 μl) was refreshed daily until 120h. When the addition of specific factors to Gastruloids was required 24h AA, 20 μl medium was carefully removed with a multichannel pipette, and 20 μl of N2B27 containing required factors at the appropriate concentration was added as previously described (Baillie-Johnson et al., 2015). Gastruloids were collected randomly at the different time-points analysed and were imaged using the Evos Cell imaging systems.

### Primordial Germ Cell-like cell (PGCLCs) differentiation

PGCLCs differentiation was performed under a floating condition as previously described (Hayashi and Saitou, 2013). Briefly, 2.5×10^4^ cells/well were plated in U-shaped ultra-low attachment 96- multiwell (Corning Costar) in a serum-free GK15 medium containing Glasgow minimal essential medium (GMEM) supplemented with 15% KSR, 0.1 mM NEAA, 1 mM sodium pyruvate, 0.1 mM 2-mercaptoethanol, 100 U/ml penicillin, 0.1 mg/ml streptomycin, and 2 mM L-glutamine. To induce PGCLC differentiation the medium was supplemented with growth factors/ cytokines including BMP4 (500 ng/ml; R&D Systems), LIF (1000 u/ml; Invitrogen), SCF (100 ng/ml; R&D Systems), and EGF (50 ng/ml; R&D Systems). After 4 days the cell aggregates were dissociated with Trypsin and the cells were used for further analysis.

### Flow cytometry and cell sorting

TBV2 mESCs were dissociated with accutase mix (1x for 5 min at 37°C) to obtain a single cell suspension and sorted with a FACS ARIAIII (Becton Dickinson) on the basis of Forward (FSC-A) and side scatter (SSC-A) parameters, excluding cellular debris and cells death.

### RNA extraction and quantitative real-time PCR

Total RNAs were isolated using either the RNeasy kit or Trizol reagent (Invitrogen) and reverse transcribed using QuantiTect Reverse Transcription kit (Qiagen). qPCR was performed using SYBR Green PCR master mix (FluoCycle II TM SYBR, EuroClone).

### Preparation of cytospin samples

Cells were dissociated with accutase or trypsin-EDTA for 5 min at 37°C, re-suspended in 15% FBS/1x PBS and centrifuged at 800 rpm for 8 min onto glass slides (2 spots, 5×10^5^ cells/spot) using a Thermo Shandon Cytocentrifuge (CytoSpinTM 4). Specimens were fixed with PFA 4% for further analysis.

### Immunofluorescence analysis

Gastruloids were fixed (4% PFA) and the immunofluorescence was performed as previously described (Baillie-Johnson et al., 2015). The primary antibodies, listed in Table S1, were used overnight at 4°C. After washing gastruloids were incubated with the appropriate secondary antibodies (Alexa Fluor DAR-594, 1:400, Molecular Probes #A21207; Alexa Fluor GAM-488, 1:400, Molecular Probes #A11001, Alexa Fluor DAG-594 1:400, Molecular Probes #). Cell nuclei were counterstained with DAPI (Invitrogen). Cytospin cell samples were fixed in 4% PFA and permeabilized (0.1% Triton X-100) for 10 minutes at RT and incubated with the primary antibodies, listed in Table S1, overnight at 4°C. After washing in 0,5% Tween-1x PBS, cells were incubated with the appropriate secondary antibodies (see above). Cell nuclei were counterstained with DAPI (Invitrogen). Images were obtained using the DMI6000B microscope and the DFC 350FX B/W digital camera (Leica Microsystems). Confocal images were obtained on a Nikon A1 microscope. The AF6000 (Leica Microsystems) and NIS Element C (Nikon, Tokyo) software were used for image acquisition/elaboration.

### Histological analysis

Gastruloid were fixed in 2% glutaraldehyde/4% paraformaldehyde, post-fixed in osmium tetraoxide, dehydrated, and embedded in Epon 812 (Polyscience, Niles, IL, USA). Thin (5 μm) sections were cut using a Leica ultracut UCT ultramicrotome (Leica Microsystems) and stained with toluidine blue (1% in water) for 10 min at RT. Images were obtained with ECLIPSE Ni-E microscope.

### Statistical analysis

Statistical significance was determined by a two-tailed paired Student’s t test. P-values of ≤0.01 were considered as statistically significant. Error bars in the figures represent SEM or SD as indicated.

## Acknowledgments

We are grateful to members of the Integrated Microscopy and FACS Facilities of IGB-ABT, CNR. We thank Gennaro Andolfi for excellent technical assistance. We are grateful to Maurizio Iaccarino; Annalisa Fico and Antonio Baldini for helpful discussions.

## Author contributions

F.C., A.M-A., G.M., and E.J.P. conceived and designed the study. F.C., C.D’A., R.T., D.D.D., and E.J.P. performed the experiments. F.C. performed statistical analysis. C.D’A., A.M-A., D.D.D., and G.M. gave conceptual advice and contribute to the editing of the manuscript. F.C. and E.J.P. wrote the manuscript.

